# OpenLipid: a large language model workflow for targeted analysis of DIA mass spectrometry data in lipidomics

**DOI:** 10.64898/2026.09.12.751192

**Authors:** Jiayi Li, Hannes Röst

**Affiliations:** Donnelly Centre for Cellular and Biomolecular Research, University of Toronto, Toronto, Ontario, Canada

## Abstract

Mass spectrometry has become a central technology for lipidomics, with data-independent acquisition (DIA) enabling broad and reproducible sampling of lipid signals. However, the multiplexed fragment-ion spectra in DIA data complicate lipid identification. Here, we introduce OpenLipid, a large language model (LLM)-based workflow for targeted DIA lipidomics. Using assay libraries built from data-dependent acquisition (DDA) results, OpenLipid directly evaluates extracted ion chromatograms (XICs) from DIA data in a zero-shot setting to identify target lipid peaks and generate human-readable rationales for individual lipid identification decisions. We benchmarked OpenLipid against manual annotations across four datasets comprising human plasma and mouse feces analyzed in positive and negative ionization modes. The plasma assay libraries contained 199 target lipids in positive mode and 147 in negative mode. The fecal assay libraries contained 181 target lipids in positive mode and 264 in negative mode. At a 5% false discovery rate (FDR) threshold, OpenLipid identified 110 (55.3%) and 28 (19.0%) library targets in plasma and 130 (71.8%) and 84 (31.8%) in feces, in positive and negative ionization modes, respectively. OpenLipid achieved an overall identification rate comparable to that of DIAMetAlyzer (57.8%, 19.7%, 89.0%, and 17.8% across the corresponding datasets) and substantially higher than that of untargeted MS-DIAL DIA analysis (12.6%, 0.0%, 47.5%, and 0.0%) on the same assay-library targets. LLM-derived chromatographic features also enabled supervised discrimination between correct and incorrect candidate peak groups for target lipids, with median cross-validation average precision values of 0.838-0.912. Together, these results demonstrate that OpenLipid is an effective LLM-based workflow for FDR-controlled targeted analysis of DIA lipidomics data.

## Introduction

Lipids are essential components of biological systems, serving as structural constituents of cellular membranes, energy stores, and signaling molecules that regulate diverse cellular processes^1,2^. Their structural diversity and varied abundance contribute to the complexity of biological lipidomes, as illustrated by the extensive range of lipid species identified in human plasma^3^. Lipidomics, the systematic characterization of lipids within cells, tissues, or organisms, provides a method to investigate how lipid composition and metabolism relate to cellular function, physiological adaptation, and disease^4,5^. Such analyses are typically enabled by mass spectrometry, particularly liquid chromatography coupled with tandem mass spectrometry (LC-MS/MS), which supports large-scale identification and quantification of lipids in complex biological samples^6^. Advances in MS-based measurements and computational analysis have further expanded lipidome coverage, providing opportunities to investigate lipid diversity across biological systems^7–9^.

Two widely used acquisition strategies in LC-MS/MS-based lipidomics are data-dependent acquisition (DDA) and data-independent acquisition (DIA). DDA selects precursor ions for fragmentation based on survey-scan information, typically prioritizing abundant ions, whereas DIA systematically fragments ions within predefined precursor isolation windows and can provide broader analyte coverage and more consistent detection across runs^10–15^. Despite these advantages, DIA data remain challenging to analyze because the multiplexed spectra contain fragments from multiple co-isolated precursors. Untargeted DIA analysis tools such as MS-DIAL^7–9^ use chromatographic deconvolution to reconstruct MS/MS spectra for subsequent spectral-library matching and lipid identification. However, chromatographic deconvolution can struggle to separate contributions from precursors with highly similar elution profiles, complicating interpretation of the resulting spectra^16^. An alternative strategy is spectral-library-guided targeted analysis of DIA data, which was established in proteomics through targeted data extraction and its automated implementation in OpenSWATH^11,17,18^, and implemented in metabolomics through DIAMetAlyzer^19^. In this strategy, DDA-derived assay libraries can provide precursor, fragmention, and retention-time information to guide chromatogram extraction and evaluation of predefined targets in DIA data. Determining whether a lipid target is detected then requires evaluating competing chromatographic peaks and distinguishing target signals from interference. For example, researchers can use Skyline to manually inspect and evaluate chromatographic evidence in targeted small-molecule and lipid analysis^20–22^. However, manual review of individual lipid targets is labor-intensive and can be subjective, limiting its scalability for large-scale studies. Thus, automated evaluation of chromatographic evidence is important for scalable targeted DIA lipidomics, supporting reliable lipid identification and quantification.

Large language models (LLMs)^23–26^ offer an opportunity to approach chromatographic evaluation through natural-language instructions that define analytical objectives and expert assessment criteria. In our previous study, ChatDIA demonstrated the feasibility of applying general-purpose LLMs to targeted DIA proteomics analysis without mass spectrometry-specific training or fine-tuning^27^. ChatDIA directly evaluated extracted ion chromatograms (XICs) using evidence such as retention-time agreement, fragment-ion co-elution, and consistency with spectral-library information to select analyte peak groups and generate confidence estimates and human-readable rationales. The framework supported both automated analysis through an application programming interface and interactive inspection through natural language, allowing researchers to examine individual decisions alongside the underlying chromatographic evidence. These findings motivate extending LLM-based chromatographic evaluation to lipidomics and systematically assessing its ability to identify target lipid signals.

Here, we introduce OpenLipid, a targeted DIA lipidomics workflow that builds assay libraries from DDA results and uses LLMs to evaluate XICs for lipid identification. OpenLipid evaluates XICs from DIA data to select target lipid peaks and generate candidate scores and individual chromatographic feature assessments. We evaluated the workflow against manual annotations across four benchmark datasets comprising human plasma and mouse feces analyzed in positive and negative ionization modes, and compared its results with those from untargeted DIA analysis using MS-DIAL and targeted DIA analysis using DIAMetAlyzer. At 5% empirical FDR, OpenLipid identified 110 of 199 library targets (55.3%) in positive mode and 28 of 147 (19.0%) in negative mode for plasma, and 130 of 181 (71.8%) in positive mode and 84 of 264 (31.8%) in negative mode for feces. The overall identification rate was comparable to that of DIAMetAlyzer, which achieved 57.8%, 19.7%, 89.0%, and 17.8% across the four datasets, and substantially higher than that of untargeted MS-DIAL DIA analysis, which achieved 12.6%, 0.0%, 47.5%, and 0.0% on the same assay-library targets. Furthermore, classifiers trained on LLM-derived features to distinguish correct from incorrect candidate peak groups achieved median average precision values of 0.838-0.912 in within-dataset cross-validation and retained predictive information during transfer between plasma and feces. Together, these results demonstrate that OpenLipid provides an effective LLM-based strategy for FDR-controlled targeted analysis of DIA lipidomics data while generating human-readable rationales that allow researchers to examine individual lipid identification decisions.

## Materials and Methods

### Benchmarking datasets and manual annotations

Public lipidomics data for NIST SRM 1950 human plasma and mouse feces were obtained from the RIKEN DROPMet repository under accession DM0050^28^. The data were acquired using the SCIEX ZenoTOF 7600 mass spectrometry platform. Four benchmarking datasets were defined according to sample type and ionization polarity: SRM 1950 plasma in positive ion mode, SRM 1950 plasma in negative ion mode, feces in positive ion mode, and feces in negative ion mode. For each sample type and ionization mode, five SWATH-DIA files covering distinct precursor m/z acquisition ranges were used. Sample-specific lipid identifications were obtained from MS-DIAL alignment analyses of six SRM 1950 plasma DDA runs and five fecal DDA runs acquired from the corresponding samples^28^. Untargeted DIA lipid identifications used for comparison were obtained from the corresponding MS-DIAL SWATH-DIA analyses^28^.

Following assay-library construction, the four datasets contained 199, 147, 181, and 264 lipid targets, respectively. Target presence and retention-time intervals were manually annotated from the XICs. This yielded 170 present and 29 absent targets for SRM 1950 plasma in positive ion mode, 75 present and 72 absent targets for SRM 1950 plasma in negative ion mode, 169 present and 12 absent targets for feces in positive ion mode, and 160 present and 104 absent targets for feces in negative ion mode.

### Spectral library construction, DIA data processing, and chromatogram extraction

Vendor-format DIA files were converted to mzML files using ProteoWizard MSConvert (v3.0.25279)^29^. Sample- and polarity-specific assay libraries were constructed from the corresponding MS-DIAL DDA analysis results^28^. For each lipid, precursor m/z, retention time, adduct, charge state, and MS/MS fragment peaks were obtained. Up to five of the most intense product ions were retained, with a minimum separation of 50 ppm between product-ion m/z values. Lipids with fewer than two retained product-ion transitions were excluded. The resulting assays were filtered according to the precursor m/z acquisition range of each SWATH-DIA file. The original DDA results contained 199, 147, 181, and 267 lipid targets for SRM 1950 positive mode (Plasma POS), SRM 1950 negative mode (Plasma NEG), feces positive mode (Feces POS), and feces negative mode (Feces NEG), respectively, of which 199, 147, 181, and 264 were retained in the corresponding assay libraries.

DIA chromatograms were extracted using DIAMetAlyzer^19^, which is built on OpenSWATH (v3.0.0-pre-develop)^18^. The DIAMetAlyzer workflow also performed chromatographic peak picking and peak-group scoring with default parameter settings. These peak-group scores were used only for DIAMetAlyzer benchmarking and were not provided to OpenLipid.

### Model configuration, prompting, and output

OpenLipid was implemented using the OpenAI GPT-5.6 Luna model accessed through the OpenAI API, with reasoning effort set to high. No external tools were enabled during inference. Each lipid target was analyzed independently, and three replicate runs (R0, R1, and R2) were performed using identical inputs, prompts, and model configurations.

For each target, the model was provided with assay-library metadata and the corresponding precursor and product-ion XICs represented as numerical RT-intensity arrays, following the representation used in ChatDIA^27^. DIAMetAlyzer-derived candidate positions, scores, rankings, and confidence estimates, as well as manual annotations, were not provided to the model. The prompt instructed the model to identify plausible peak-group candidates directly from the ion traces, estimate their apex retention times and peak boundaries, and evaluate them according to retention-time agreement, ion co-elution, chromatographic peak shape, and fragment-ion pattern consistency. The model then selected the best-supported candidate or reported that no convincing target lipid peak was present.

### Evaluation metrics and statistical analysis

Manual annotations were used as the ground truth. For lipid-present targets, a selected peak group was considered correct when its retention-time interval matched the manually annotated interval. For lipid-absent targets, a no-peak decision was considered correct. Decision accuracy was calculated as the proportion of correct peak/no-peak decisions across all annotated targets.

For OpenLipid and DIAMetAlyzer, FDR-coverage and precision-recall were evaluated based on the best candidate peak groups. OpenLipid candidates were ranked using the LLM-derived score, whereas DIAMetAlyzer candidates were ranked using the peak-group score. These analyses used the best available candidate independently of the final peak/no-peak decision. For MS-DIAL^9^ DIA evaluation, only lipid identifications corresponding to assay-library targets were considered for comparison. The MS-DIAL-reported “Total score” was used to define score thresholds for FDR analysis. An MS-DIAL lipid identification was considered correct when its reported RT matched the manual annotation and incorrect otherwise. Unmatched assay-library targets remained in the coverage denominator but could not be accepted. For all three methods, FDR was defined as the proportion of accepted identifications that were incorrect, whereas coverage was defined as the proportion of annotated library targets that were accepted. Standard precision-recall analyses were also performed.

The LLM replicates were evaluated individually and combined using Conservative, Majority, and Open ensembles, with peaks matched by overlapping predicted retention-time intervals. Conservative required support from all replicates and used intersecting boundaries and mean scores; Majority required support from at least half of the replicates and used median boundaries and mean scores weighted by replicate support; Open retained the boundaries and score of the highest-scoring replicate. For FDR-coverage and precision-recall analyses, the ensemble rules were applied to the best candidate reported by each replicate, regardless of whether the candidate was selected in the final decision. For decision accuracy, the ensemble rules were applied only to candidates selected in the final decisions, with a no-peak decision contributing no candidate.

### Supervised learning evaluation of feature representations

Separate candidate-level classifiers were trained to evaluate whether DIAMetAlyzer- and LLM-derived feature representations could distinguish correct from incorrect peak-group candidates. A candidate was labeled as correct when its retention-time interval matched the manually annotated peak and incorrect otherwise; all candidates from lipid-absent targets were labeled as incorrect. Candidates with invalid retention-time boundaries were excluded. DIAMetAlyzer models used native numerical features describing library similarity, ion co-elution, chromatographic peak shape, mass accuracy, isotope patterns, signal intensity, signal-to-noise ratio, and retention-time deviation. LLM models used the LLM-reported scores for retention-time agreement, ion co-elution, chromatographic peak shape, and fragment-ion pattern consistency, together with the number of supporting product ions. Native aggregate or discriminant scores, final selection outputs, identifiers, and raw retention-time coordinates were excluded.

L2-regularized logistic regression models were trained for the two feature representations using the same model settings. Performance was evaluated using repeated stratified five-fold cross-validation with 10 repeats. Data were split by lipid target and stratified by the manually annotated lipid-present/absent status, ensuring that candidates from the same target did not occur in both training and test folds. Individual LLM runs were modeled separately.

Candidate-level performance was evaluated using precision-recall analysis and primarily summarized using average precision. Among lipid targets for which the candidate set included the correct peak-group candidate, top-1 accuracy was defined as the proportion for which a correct candidate received the highest predicted probability. Feature importance was summarized as the mean absolute logistic regression coefficient for each standardized input feature across cross-validation models. External generalization was evaluated bidirectionally between plasma and feces within the same ionization polarity, with all preprocessing and feature filtering determined from the training data. Probability cutoffs corresponding to 1% and 5% training FDR were derived from out-of-fold predictions and applied unchanged to the corresponding external test datasets. Because DIAMetAlyzer and the LLM proposed different candidate sets, the analysis evaluated the predictive information within each feature representation rather than constituting a fully matched head-to-head comparison.

## Results

### OpenLipid enables automated targeted DIA lipid identification

We developed OpenLipid, an LLM-based workflow for automated targeted analysis of DIA lipidomics data (Figure 1). Sample-specific DDA data were first used for lipid identification (by MS-DIAL) and spectral-library construction. The resulting library guided the extraction of precursor and product-ion XICs from the corresponding DIA data. These chromatographic signals were then analyzed by LLMs within OpenLipid.

**Figure 1.**
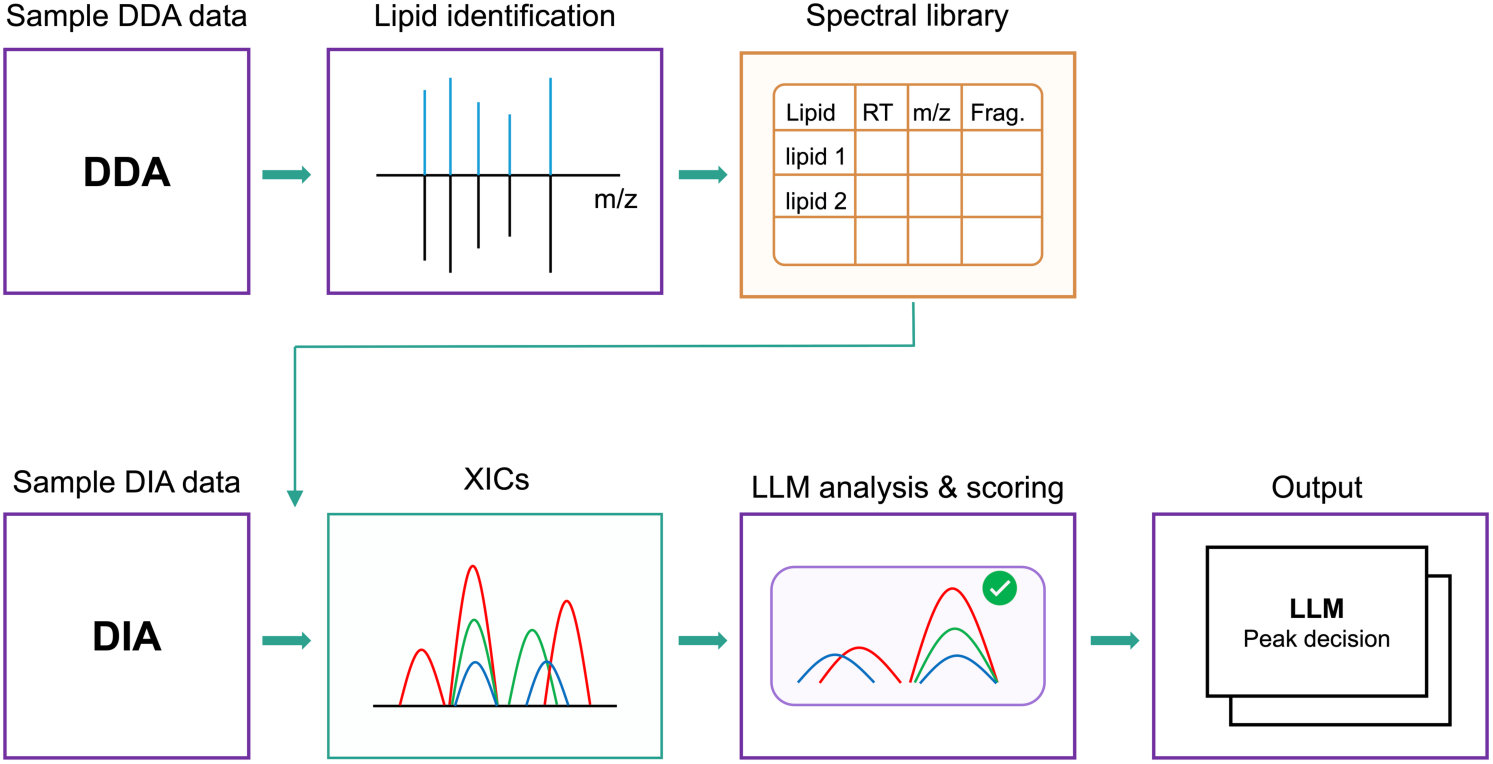
Overview of the OpenLipid workflow for targeted DIA lipidomics. Sample-specific data-dependent acquisition (DDA) data were used for lipid identification and spectral library construction. The resulting library guided the extraction of ion chromatograms (XICs) from corresponding data-independent acquisition (DIA) data. The large language model (LLM)-based OpenLipid workflow directly identified and evaluated candidate peak groups from the XICs, producing candidate scores and peak-selection decisions.

For each lipid target, OpenLipid uses assay-library metadata and numerical XIC arrays to identify plausible peak groups directly from ion traces. The model assesses retention-time agreement, ion co-elution, peak shape, and fragment-ion pattern consistency with the assay library to select the best-supported peak or report no convincing target peak. OpenLipid produces structured candidate scores and a final peak/no-peak decision with peak boundaries and supporting product ions. This chromatographic evaluation is performed by general-purpose LLMs without access to DIAMetAlyzer-derived candidate positions or scores and without lipidomics-specific training or fine-tuning, enabling automated targeted analysis of DIA lipidomics data.

### OpenLipid provides broad lipid identification coverage across diverse benchmark datasets

We evaluated OpenLipid using SRM 1950 plasma and mouse feces datasets acquired in positive and negative ionization modes (Figure 2). MS-DIAL identified 199, 147, 181, and 267 unique lipids from DDA data and 198, 152, 165, and 296 from DIA data in Plasma POS, Plasma NEG, Feces POS, and Feces NEG, respectively. Only 93, 88, 103, and 160 lipid identities were shared between the corresponding DDA and DIA results, indicating substantial differences in the lipids identified by MS-DIAL between the two acquisition modes (Figure 2a). Manual inspection confirmed DIA lipid presence for 170 of 199 DDA-derived targets in Plasma POS, 75 of 147 in Plasma NEG, 169 of 181 in Feces POS, and 160 of 264 in Feces NEG (Figure 2b).

**Figure 2.**
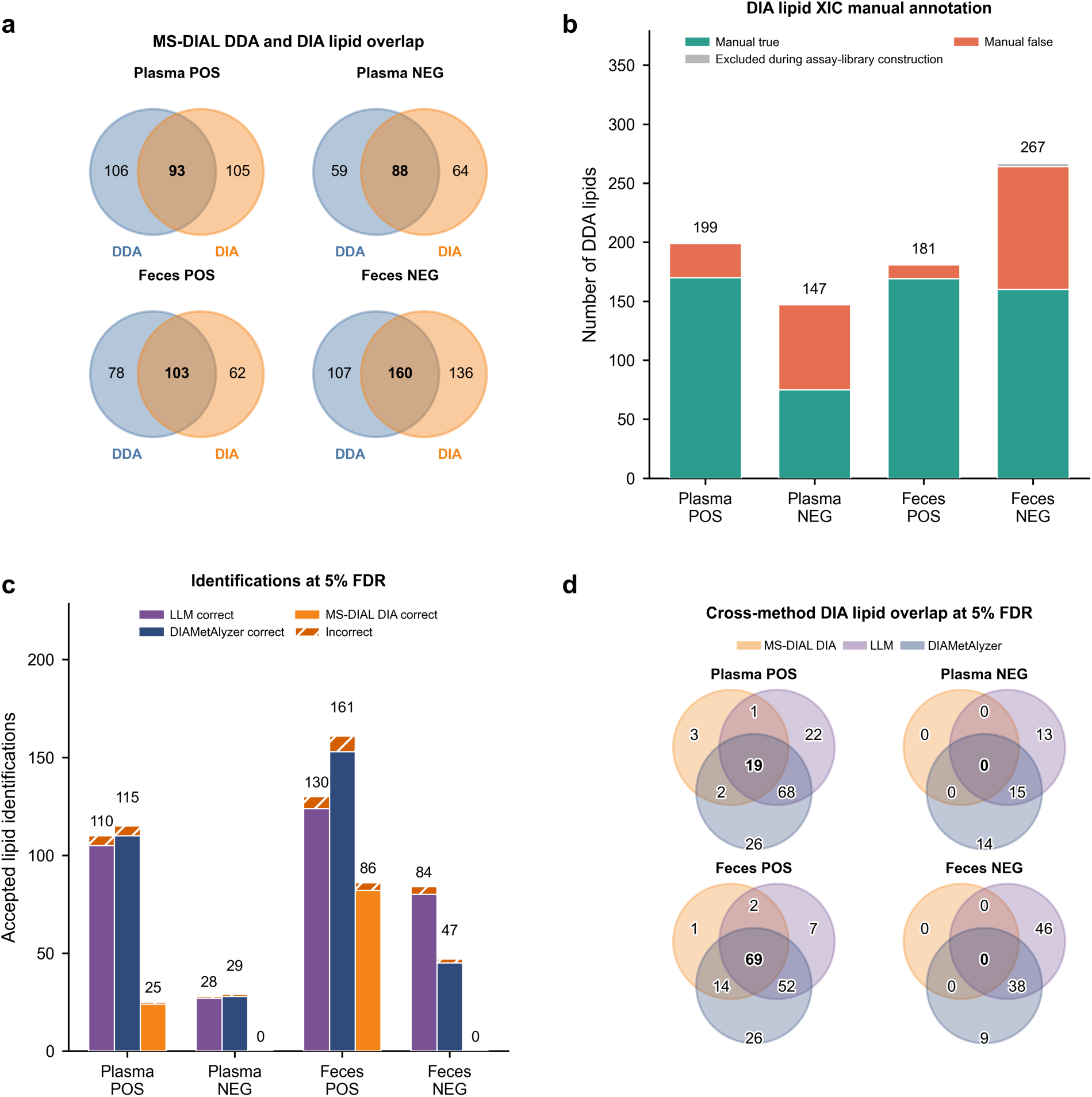
Overall comparison of DDA and DIA lipid identifications. Mouse feces and Standard Reference Material 1950 (SRM 1950) plasma were analyzed in positive (POS) and negative (NEG) ionization modes. (a) Overlap between unique lipid identities reported by MS-DIAL DDA and DIA analyses. (b) Manual XIC annotation of DDA-derived lipid assay targets in the DIA data, classified as manual true, manual false, or excluded during assay-library construction. (c) Correct and incorrect lipid target identifications accepted by the OpenLipid LLM Majority ensemble, DIAMetAlyzer, and MS-DIAL DIA at method-specific score thresholds that maximized coverage while maintaining an empirical false discovery rate (FDR) of no more than 5%. (d) Dataset-specific overlap among lipid identities accepted by MS-DIAL DIA, OpenLipid LLM Majority, and DIAMetAlyzer at 5% FDR. In (c) and (d), MS-DIAL DIA was evaluated using the same assay-library target lipids as OpenLipid LLM Majority and DIAMetAlyzer. In (a) and (d), lipid identity matching required identical polarity, SMILES, and adduct, with a precursor m/z difference of no more than 25 ppm.

At 5% FDR, OpenLipid (LLM Majority) accepted 110, 28, 130, and 84 identifications in Plasma POS, Plasma NEG, Feces POS, and Feces NEG, respectively, corresponding to 55.3%, 19.0%, 71.8%, and 31.8% of the library targets (Figure 2c). Of these accepted identifications, 105, 27, 124, and 80 were correct, respectively. DIAMetAlyzer accepted 115, 29, 161, and 47 identifications, corresponding to 57.8%, 19.7%, 89.0%, and 17.8% of the library targets, of which 110, 28, 153, and 45 were correct, respectively. At the same threshold, untargeted MS-DIAL DIA analysis accepted 25 and 86 identifications in Plasma POS and Feces POS, corresponding to 12.6% and 47.5% of the library targets, of which 24 and 82 were correct. No MS-DIAL DIA identifications satisfied the 5% FDR threshold in Plasma NEG or Feces NEG. OpenLipid therefore achieved overall coverage comparable to that of DIAMetAlyzer and substantially higher than that of MS-DIAL DIA analysis. Across all three methods, coverage varied among datasets and was consistently higher in positive than in negative ionization mode.

The identification sets produced by OpenLipid and DIAMetAlyzer overlapped substantially, with greater overlap in positive than in negative ionization mode (Figure 2d). Each targeted method also contributed identifications not recovered by the other, particularly OpenLipid in Feces NEG and DIAMetAlyzer in Feces POS. Among assay-library targets, nearly all MS-DIAL DIA identifications accepted at 5% FDR in positive ionization mode overlapped with one or both targeted approaches. Together, these results show that OpenLipid and DIAMetAlyzer recovered broader sets of assay-library targets than untargeted MS-DIAL DIA analysis.

### OpenLipid supports lipid identification at low FDR

We next compared the ability of OpenLipid, DIAMetAlyzer, and MS-DIAL DIA scores to prioritize correct identifications under FDR control (Figure 3, Supplementary Figure 1, and Supplementary Table 1). At a 1% FDR threshold, OpenLipid achieved coverage of 35.7%, 12.2%, 39.2%, and 22.7% in Plasma POS, Plasma NEG, Feces POS, and Feces NEG, respectively, with no incorrect candidates observed among the accepted identifications. The corresponding coverage was 33.7%, 4.8%, 70.7%, and 15.5% for DIAMetAlyzer and 6.0%, 0.0%, 16.6%, and 0.0% for MS-DIAL DIA. OpenLipid achieved higher coverage than DIAMetAlyzer in three of the four datasets at this stringent threshold, while DIAMetAlyzer achieved substantially higher coverage in Feces POS. Both targeted approaches achieved higher identification coverage than MS-DIAL DIA across all four datasets.

**Figure 3.**
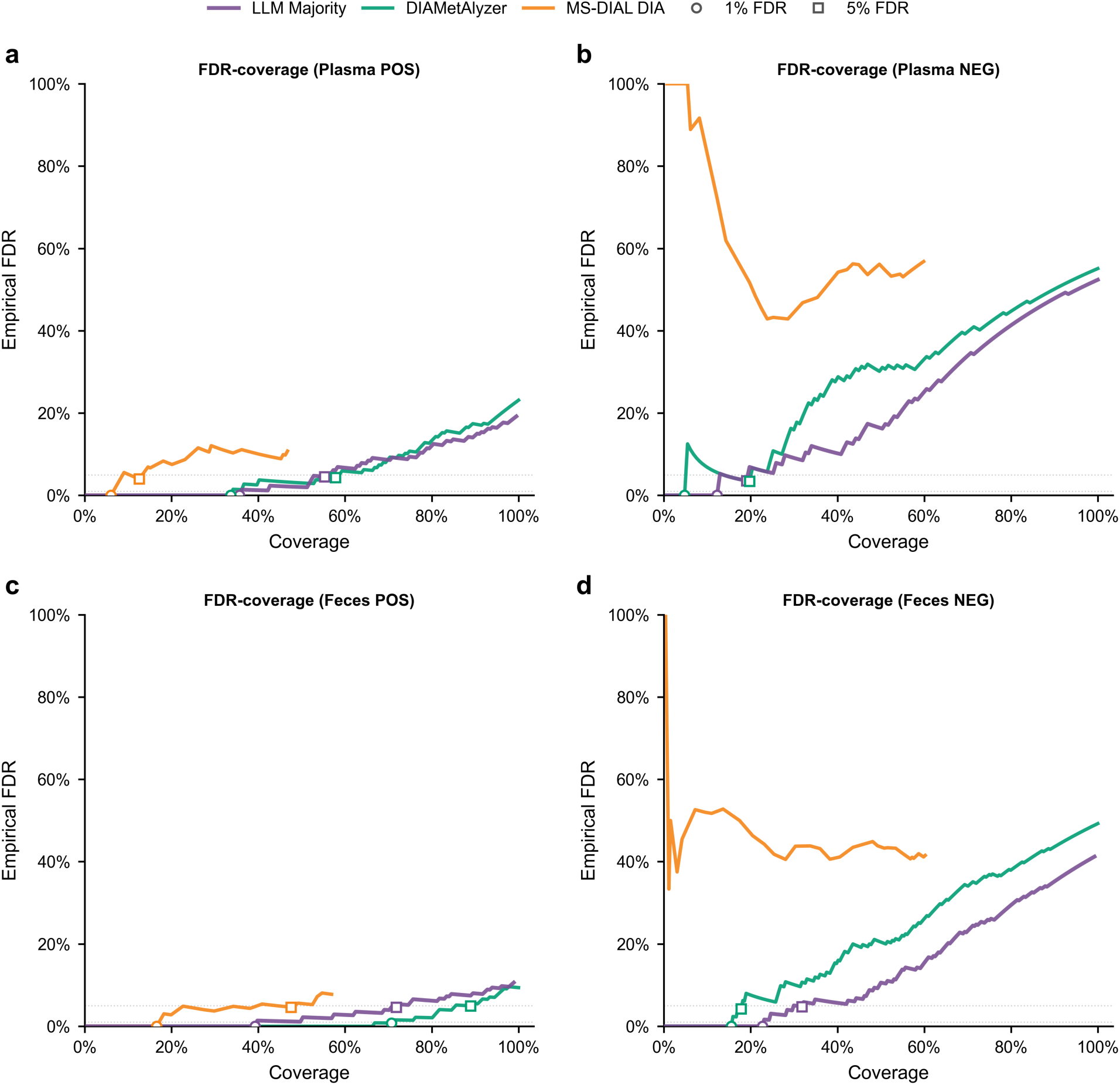
FDR-coverage analysis of DIA lipidomics methods. FDR-coverage curves for OpenLipid LLM Majority, DIAMetAlyzer, and MS-DIAL DIA are shown for (a) Plasma POS, (b) Plasma NEG, (c) Feces POS, and (d) Feces NEG. For OpenLipid, candidate-ranking analyses retained the best candidate even when the final LLM decision was no peak. Coverage was calculated as the number of accepted candidates divided by all manually annotated target lipids, and empirical FDR was the fraction of accepted candidates that were incorrect. Circles and squares indicate the maximum coverage achieved at empirical FDR ≤1% and ≤5%, respectively.

Increasing the FDR threshold from 1% to 5% increased coverage for OpenLipid and DIAMetAlyzer across all four datasets and for MS-DIAL DIA in the two positive-ionization datasets. At 5% FDR, OpenLipid and DIAMetAlyzer achieved comparable overall identification coverage, with similar coverage in both plasma datasets, higher coverage for OpenLipid in Feces NEG, and higher coverage for DIAMetAlyzer in Feces POS. Both targeted approaches achieved higher coverage than MS-DIAL DIA in every dataset.

Precision-recall analysis further compared the ability of the three methods to prioritize correct identifications (Supplementary Figure 2). OpenLipid achieved average precision values of 0.903, 0.851, 0.924, and 0.900 in Plasma POS, Plasma NEG, Feces POS, and Feces NEG, respectively, compared with 0.865, 0.732, 0.960, and 0.738 for DIAMetAlyzer and 0.451, 0.232, 0.544, and 0.325 for MS-DIAL DIA. Both targeted approaches achieved substantially higher average precision than MS-DIAL DIA across all four datasets. These results support the use of OpenLipid scores for candidate ranking and lipid identification under FDR control, although the achievable coverage varied across datasets.

### Majority ensembling improves the robustness of OpenLipid across replicate runs

We assessed the reproducibility of OpenLipid using three independent LLM analyses of the same lipid targets (Figure 4). All three runs selected the same peak or consistently reported no peak for 80.4%, 80.3%, 80.7%, and 77.3% of targets in Plasma POS, Plasma NEG, Feces POS, and Feces NEG, respectively (Figure 4a). A majority consensus was reached for 97.8%-99.6% of targets across the four datasets. All three runs produced correct final decisions for 64.4%-72.4% of targets, while at least two runs were correct for 76.5%-81.8% (Figure 4b). These results indicate that decisions for individual lipid targets were largely reproducible across independent LLM runs, with run-to-run variation concentrated in a relatively small subset of targets.

**Figure 4.**
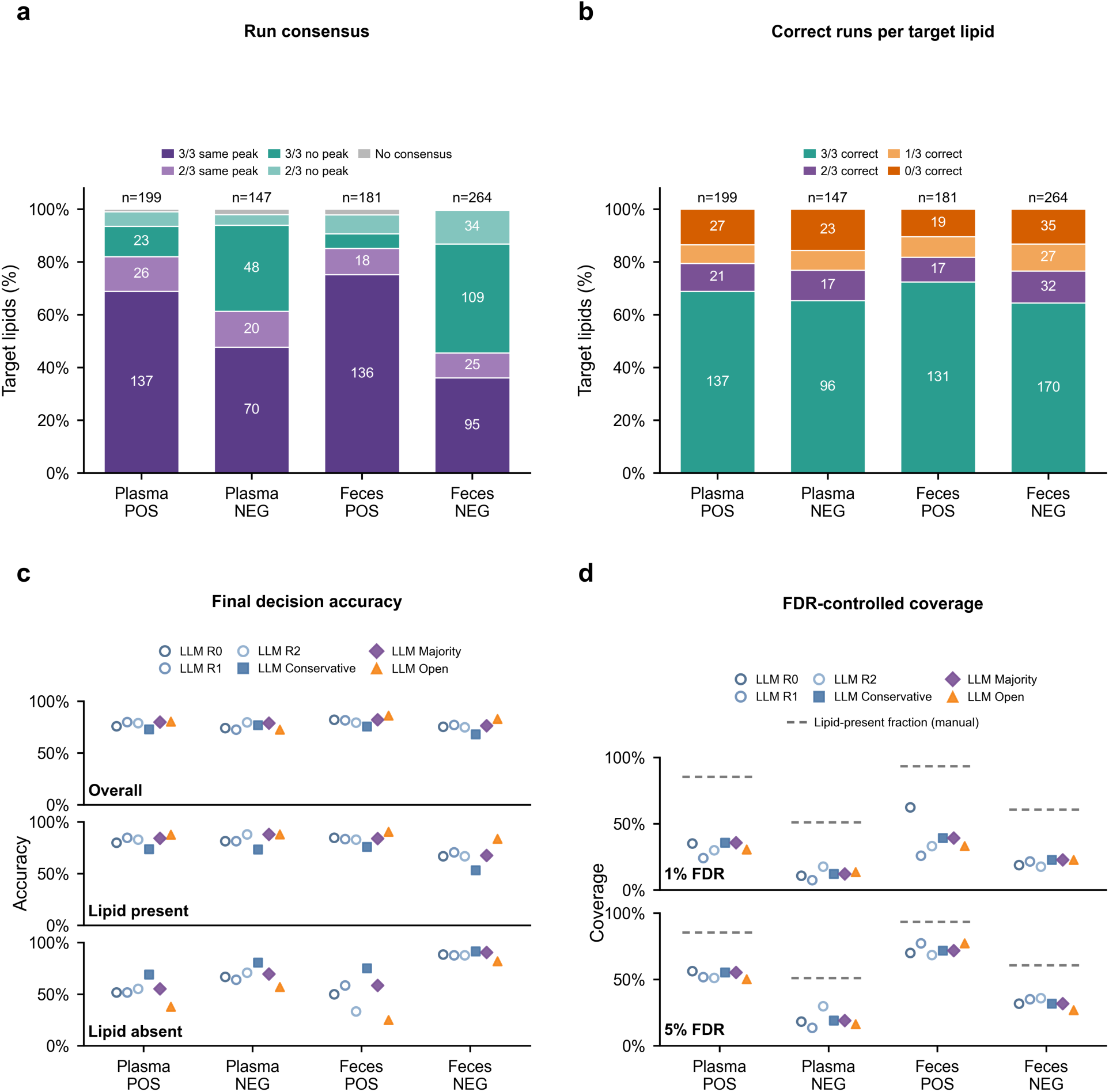
Reproducibility of independent LLM runs and comparison of ensemble decision rules. (a) Peak-selection consensus among three independent LLM runs (R0-R2). (b) Number of correct runs for each target lipid. (c) Overall, lipid-present, and lipid-absent final-decision accuracy for the individual runs and ensemble strategies. (d) Candidate coverage at empirical FDR ≤1% and ≤5%. Gray dashed segments indicate the manually annotated lipid-present fraction for each dataset. Panels (a-c) evaluate the final peak group selection decisions, whereas panel (d) evaluates candidate ranking. LLM Conservative required support from all three runs, LLM Majority required support from at least two runs, and LLM Open required support from at least one run.

The Majority ensemble achieved overall decision accuracies of 79.9%, 78.9%, 82.3%, and 76.5% in Plasma POS, Plasma NEG, Feces POS, and Feces NEG, respectively, matching or approaching the best individual-run performance in each dataset (Figure 4c). The ensemble strategies showed different decision tendencies: Conservative more frequently retained no-peak decisions and performed better for lipid-absent targets, whereas Open more readily retained predicted peaks and performed better for lipid-present targets. Majority provided a balance between these behaviors, reducing dependence on either overly conservative or permissive peak selection.

Individual-run FDR-coverage profiles also showed dataset-dependent variability (Supplementary Figure 3). For example, coverage at 1% FDR ranged from 26.0% to 62.4% among the three runs in Feces POS, while coverage at 5% FDR ranged from 13.6% to 29.9% in Plasma NEG. Applying the ensemble rules produced consensus FDR-coverage profiles across the replicate runs (Figure 4d and Supplementary Figure 4). Together, these results show that Majority ensembling reduces dependence on a single stochastic LLM run while maintaining balanced peak/no-peak decisions and FDR-controlled candidate prioritization.

### LLM-derived features support candidate classification and cross-dataset generalization

We evaluated whether the individual chromatographic features reported by the LLM could distinguish correct from incorrect peak-group candidates without using the LLM’s final evaluation score (Figure 5). Classifiers for lipid peak-group candidates were evaluated using five-fold cross-validation grouped by lipid target and repeated 10 times. The three LLM runs were modeled separately, resulting in 30 performance estimates per dataset for the LLM-derived features and 10 for the deterministic DIAMetAlyzer-derived features. Classifiers trained on LLM-derived features achieved median average precision values of 0.870, 0.838, 0.912, and 0.859 in Plasma POS, Plasma NEG, Feces POS, and Feces NEG, respectively (Figure 5a). The corresponding values for classifiers trained on DIAMetAlyzer-derived features were 0.854, 0.734, 0.865, and 0.860. All values were substantially higher than the candidate-prevalence baselines, which ranged from 0.117 to 0.270. Precision-recall curves based on predictions averaged across cross-validation repeats are shown in Supplementary Figure 5.

**Figure 5.**
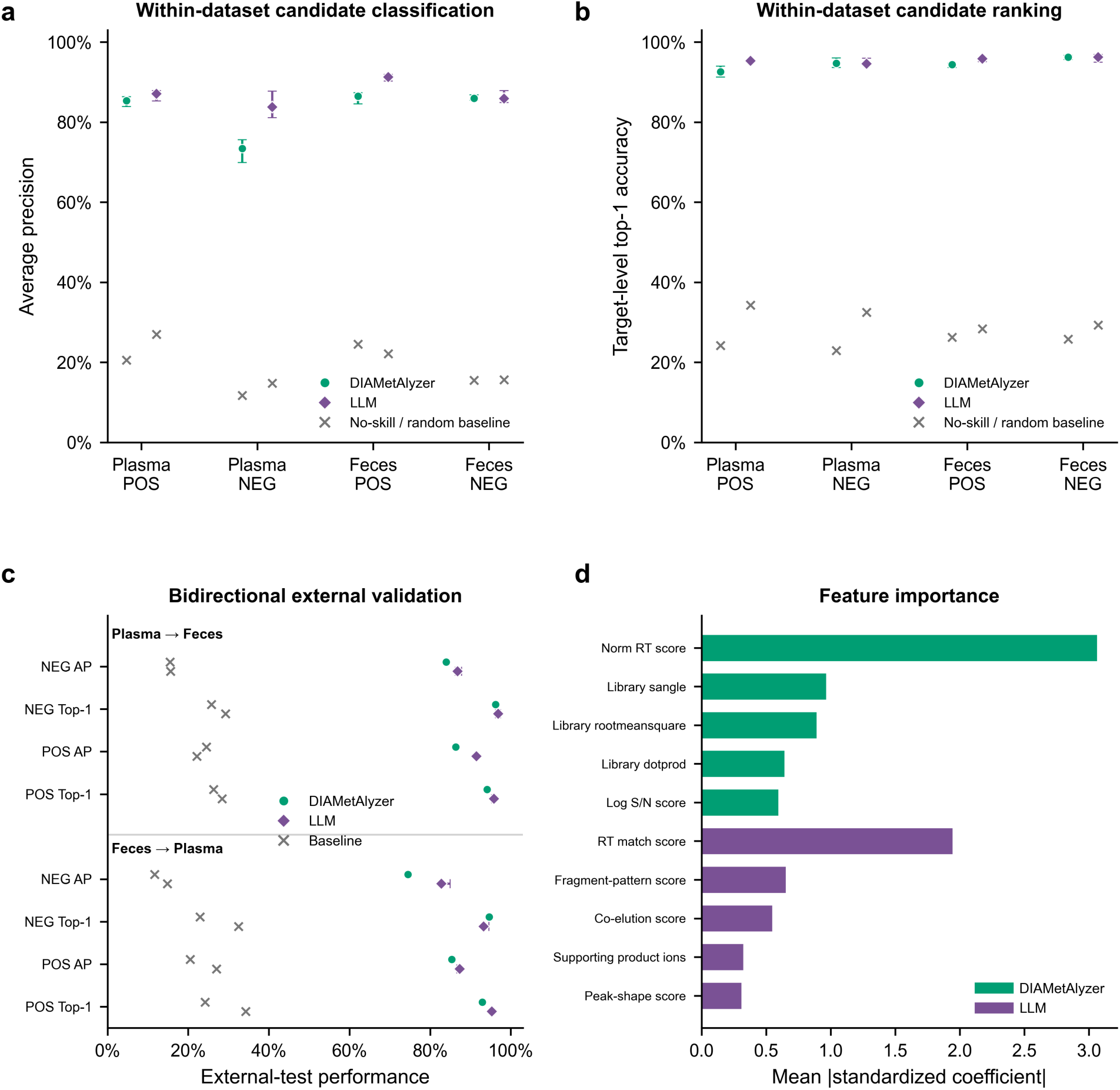
Supervised machine-learning evaluation of DIAMetAlyzer- and LLM-derived candidate features. (a) Candidate-level average precision and (b) target-level top-1 candidate-ranking accuracy under repeated target-grouped five-fold cross-validation. Points show medians and error bars show the 2.5th-97.5th percentiles across 10 repeats; LLM summaries additionally include three independent replicates (R0-R2). Gray crosses indicate candidate-prevalence or random-ranking baselines. (c) Bidirectional external validation between plasma and feces within each ionization polarity. LLM points and error bars show the median and range across R0-R2. (d) Mean absolute standardized logistic-regression coefficients for the five most influential features from each representation.

Top-1 accuracy measured how often the highest-probability candidate was correct among targets containing a correct candidate. Median top-1 accuracy ranged from 0.946 to 0.962 for classifiers trained on LLM-derived features and from 0.926 to 0.962 for those trained on DIAMetAlyzer-derived features (Figure 5b). Classifiers using LLM-derived features achieved higher top-1 accuracy in Plasma POS and Feces POS and similar accuracy in Plasma NEG and Feces NEG. These results indicate that both feature representations supported accurate candidate ranking within individual lipid targets.

The predictive information in the LLM-derived features was retained during bidirectional external validation between plasma and feces within each ionization mode (Figure 5c and Supplementary Figure 6). When models trained on plasma were applied to feces datasets, classifiers using LLM-derived features achieved median average precision values of 0.915 and 0.868 in positive and negative ionization modes, respectively, compared with 0.864 and 0.840 for those using DIAMetAlyzer-derived features. In the reverse direction, the corresponding values were 0.873 and 0.828 for LLM-derived features and 0.854 and 0.745 for DIAMetAlyzer-derived features. Median top-1 accuracy for classifiers using LLM-derived features ranged from 0.932 to 0.969 across the four external validation settings, supporting the transferability of candidate-ranking information across sample types. However, the transferability of probability cutoffs corresponding to nominal FDR levels was direction-dependent: plasma-trained models largely maintained the nominal FDR in feces, whereas feces-trained models generally exceeded the nominal FDR in plasma (Supplementary Table 2).

To assess feature importance, we examined the coefficients learned by the logistic regression classifiers used to distinguish correct from incorrect peak-group candidates, averaging their absolute values across cross-validation models fitted to standardized features (Figure 5d). Among the LLM-derived features, retention-time agreement had the largest mean absolute coefficient (1.945), followed by fragment-pattern consistency with the assay library (0.654) and co-elution (0.549). For DIAMetAlyzer-derived features, the normalized retention-time score had the largest coefficient magnitude, followed by spectral-library similarity features. Because OpenLipid and DIAMetAlyzer proposed different candidate sets, these results describe the predictive information within each feature representation rather than a fully matched comparison of the two pipelines.

## Discussion

This proof-of-concept study presents OpenLipid, a targeted DIA lipidomics workflow that integrates DDA-derived assay libraries with LLM-based chromatographic evaluation. Across plasma and fecal DIA datasets acquired in both ionization modes, the overall lipid identification rate of OpenLipid at 5% FDR was comparable to that of DIAMetAlyzer and substantially higher than that of untargeted MS-DIAL DIA analysis. These findings demonstrate that LLM-based chromatographic evaluation is an effective strategy for FDR-controlled targeted lipid identification in DIA data.

The central methodological novelty of OpenLipid is the use of LLMs to evaluate chromatographic evidence for lipid target detection and identification, extending an approach demonstrated in our previous ChatDIA study in proteomics^27^. OpenLipid analyzes XICs extracted using DIAMetAlyzer without receiving its detected peak groups or scores, allowing the LLM and conventional workflow to assess the same chromatographic data separately. Beyond its final decisions, the LLM produced individual chromatographic features that supported candidate classification and retained predictive information during transfer between plasma and feces, suggesting OpenLipid’s applicability to additional lipidomics datasets. Feature importance analysis showed that classifiers trained on either feature representation relied most strongly on retention-time agreement, with additional contributions from library-based fragment-pattern similarity and co-elution scores for OpenLipid and spectral-library similarity features for DIAMetAlyzer. These findings are consistent with the retention-time and fragment-ion evidence considered during manual chromatographic evaluation. Together, these results suggest that LLM-derived assessments can serve as informative inputs for subsequent statistical modeling, enabling downstream models to be tailored to specific analytical needs within OpenLipid-based workflows.

Reproducibility is an important consideration when using LLMs for analytical workflows. Despite run-to-run variability, most lipid targets received consistent peak selections or no-peak decisions across independent runs. As illustrated in our ChatDIA study, variation among individual runs also provides an opportunity to improve decision robustness through ensembling^27^. Here, majority ensembling maintained decision accuracy while supporting FDR-controlled candidate prioritization. Further evaluation using additional runs and repeated ensembles would help determine the number of runs needed and quantify the remaining variability.

Several limitations should be considered. First, the evaluation used four datasets acquired on a single instrument platform, so broader applicability remains to be established across laboratories, acquisition settings, and chromatographic conditions. Second, the FDR-controlled comparison with MS-DIAL DIA was restricted to assay-library targets and therefore did not evaluate MS-DIAL DIA identifications outside the library. Third, the current OpenLipid workflow relies on MS-DIAL for DDA-based lipid identification and assay-library construction, but MS-DIAL does not provide specific fragment identities in its lipid MS/MS annotations. This limits the structural information available to the LLM when evaluating fragment-ion evidence. Finally, although candidate-ranking information transferred across sample types, the transferability of score cutoffs selected at nominal FDR levels may depend on the dataset. Score cutoffs may therefore need adjustment when applied to new datasets. LLM-derived confidence scores could also help prioritize ambiguous peak groups for manual review.

More broadly, OpenLipid allows analytical objectives and expert evaluation criteria to be expressed through natural language, providing flexibility to adapt workflows to different research needs. Combining automated processing with human-readable assessments also creates opportunities for interactive review, allowing researchers to examine chromatographic evidence and question model decisions in ambiguous cases. Advances in foundation model capabilities and computational efficiency may further improve analytical performance and enable larger-scale applications. OpenLipid thus provides a starting point for integrating adaptable LLM-based approaches into DIA lipidomics workflows.

## Supplementary materials

**Supplementary Figure 1.**
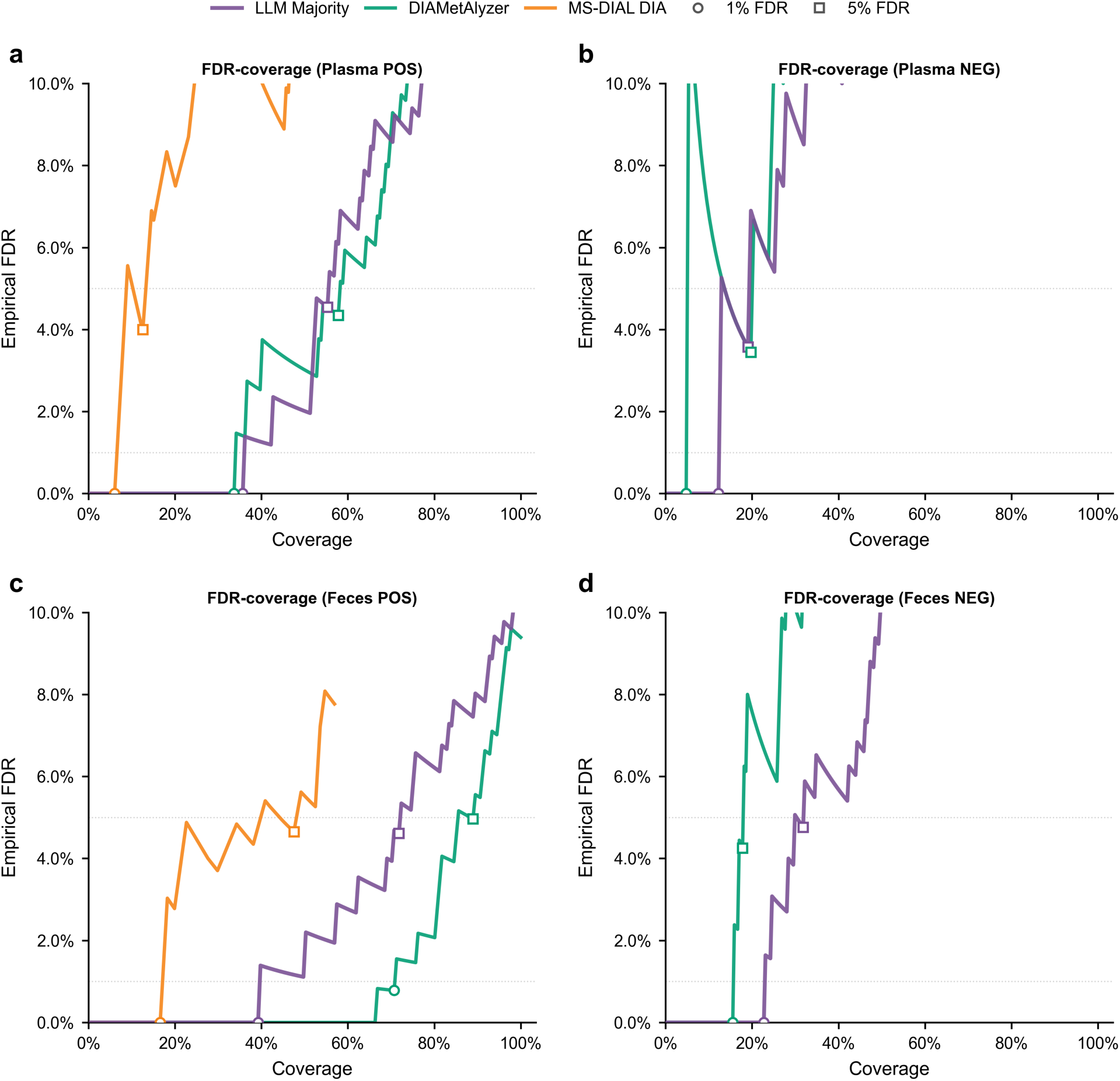
Low-FDR performance of DIA lipidomics methods. FDR-coverage curves within the 0-10% empirical FDR region for OpenLipid LLM Majority, DIAMetAlyzer, and MS-DIAL DIA are shown for (a) Plasma POS, (b) Plasma NEG, (c) Feces POS, and (d) Feces NEG. For Plasma NEG and Feces NEG, all MS-DIAL DIA operating points exceeded 10% empirical FDR and therefore fell outside the displayed range.

**Supplementary Figure 2.**
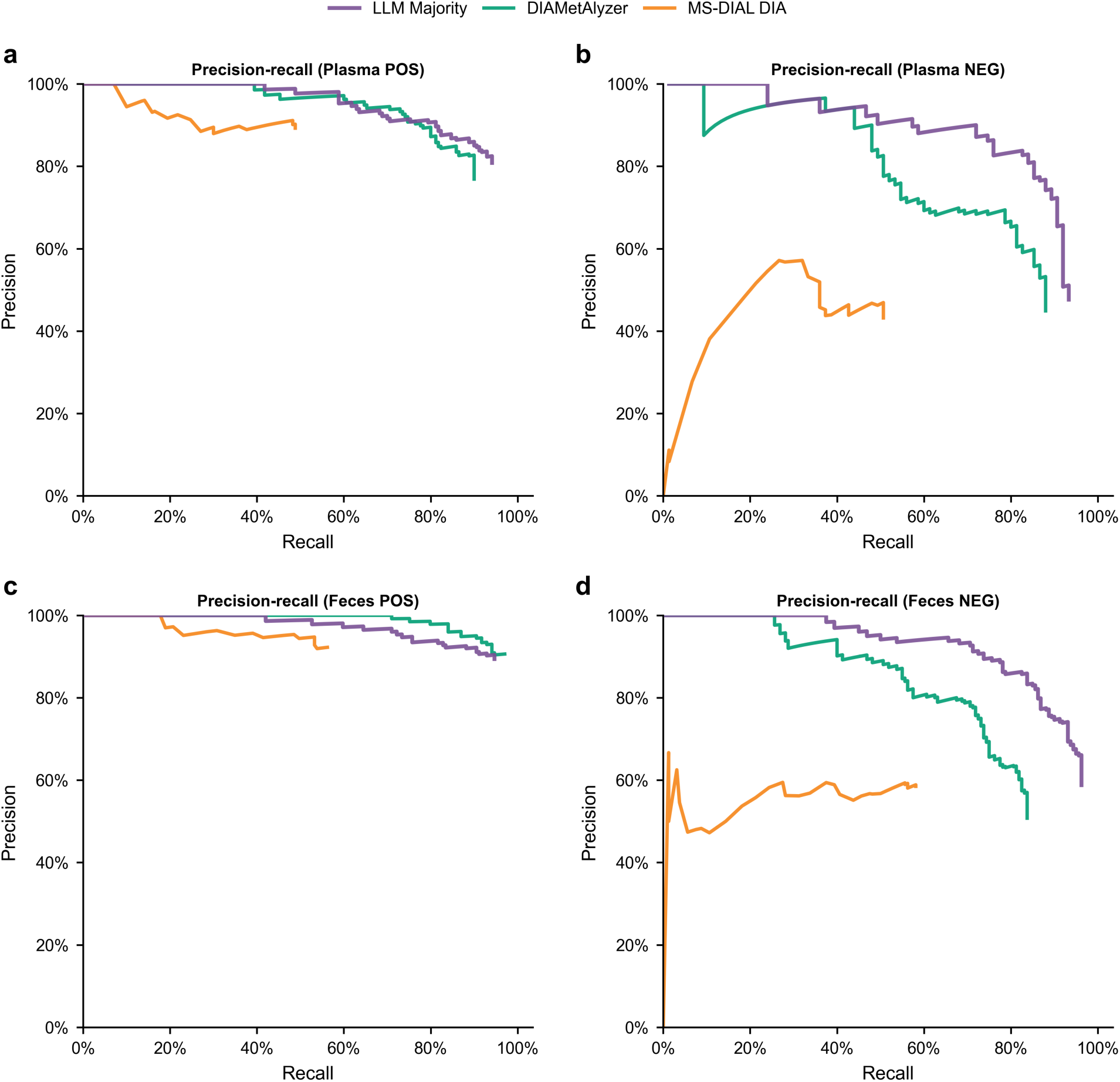
Precision-recall performance of DIA lipidomics methods. Complete precision-recall curves for OpenLipid LLM Majority, DIAMetAlyzer, and MS-DIAL DIA are shown for (a) Plasma POS, (b) Plasma NEG, (c) Feces POS, and (d) Feces NEG. Precision was the fraction of accepted candidates that were correct, and recall was calculated relative to all manually annotated lipid-present targets.

**Supplementary Figure 3.**
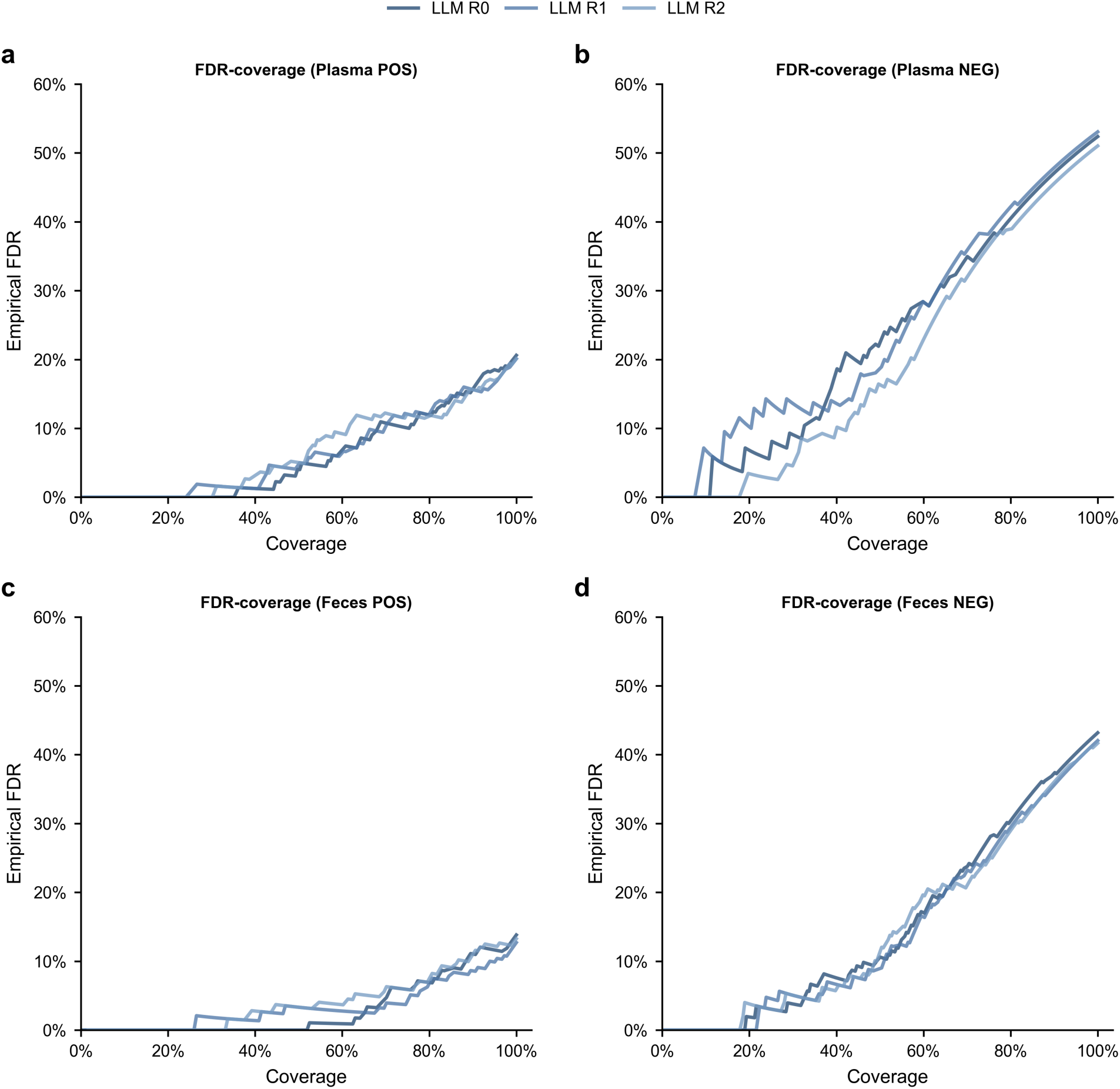
FDR-coverage profiles of independent LLM runs. Complete FDR-coverage curves for three independent LLM analyses (LLM R0, LLM R1, and LLM R2) are shown for (a) Plasma POS, (b) Plasma NEG, (c) Feces POS, and (d) Feces NEG. The runs represent repeated analyses of the same target lipids rather than biological replicates.

**Supplementary Figure 4.**
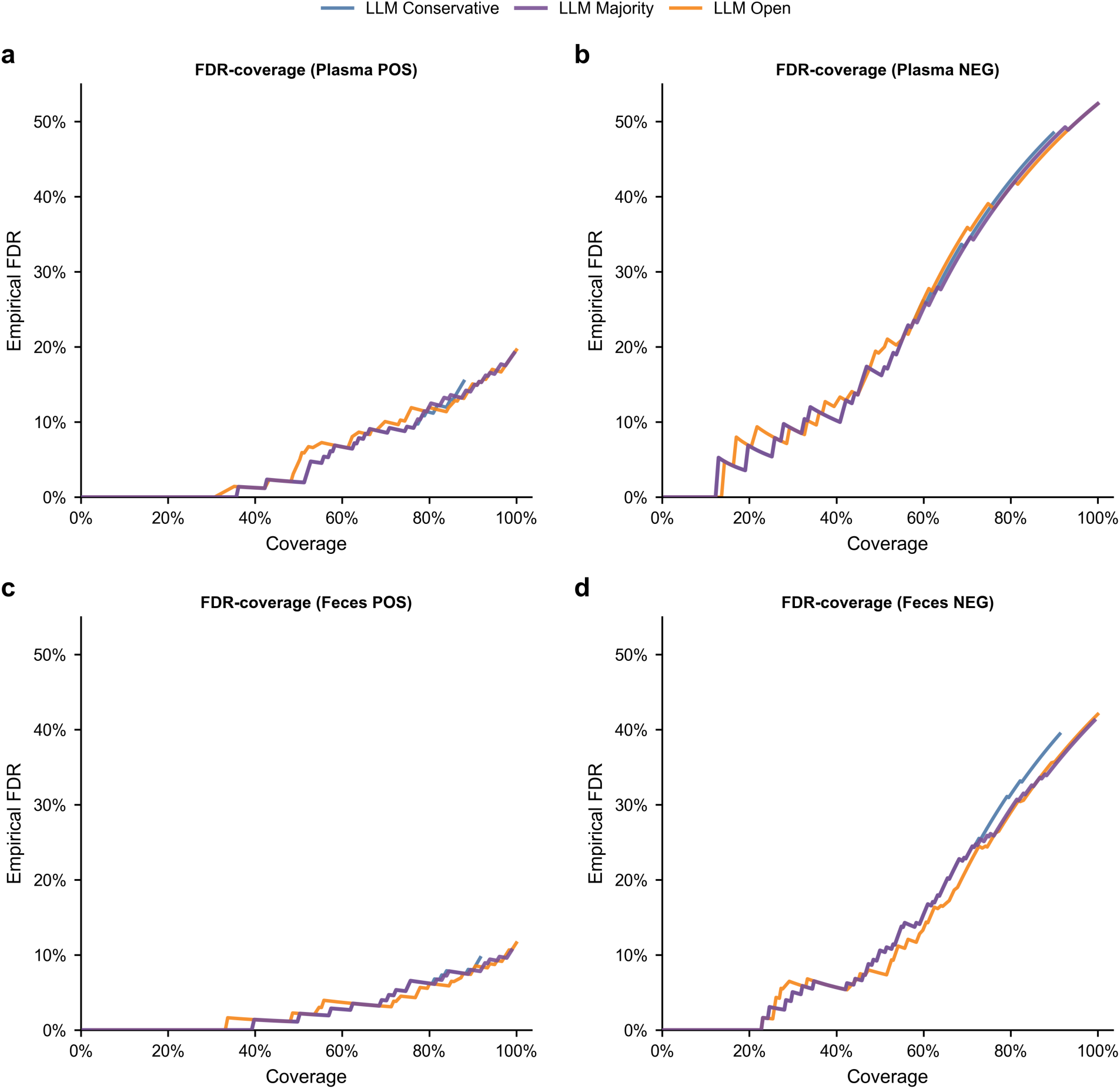
FDR-coverage profiles of LLM ensemble strategies. Complete FDR-coverage curves for LLM Conservative, LLM Majority, and LLM Open are shown for (a) Plasma POS, (b) Plasma NEG, (c) Feces POS, and (d) Feces NEG.

**Supplementary Figure 5.**
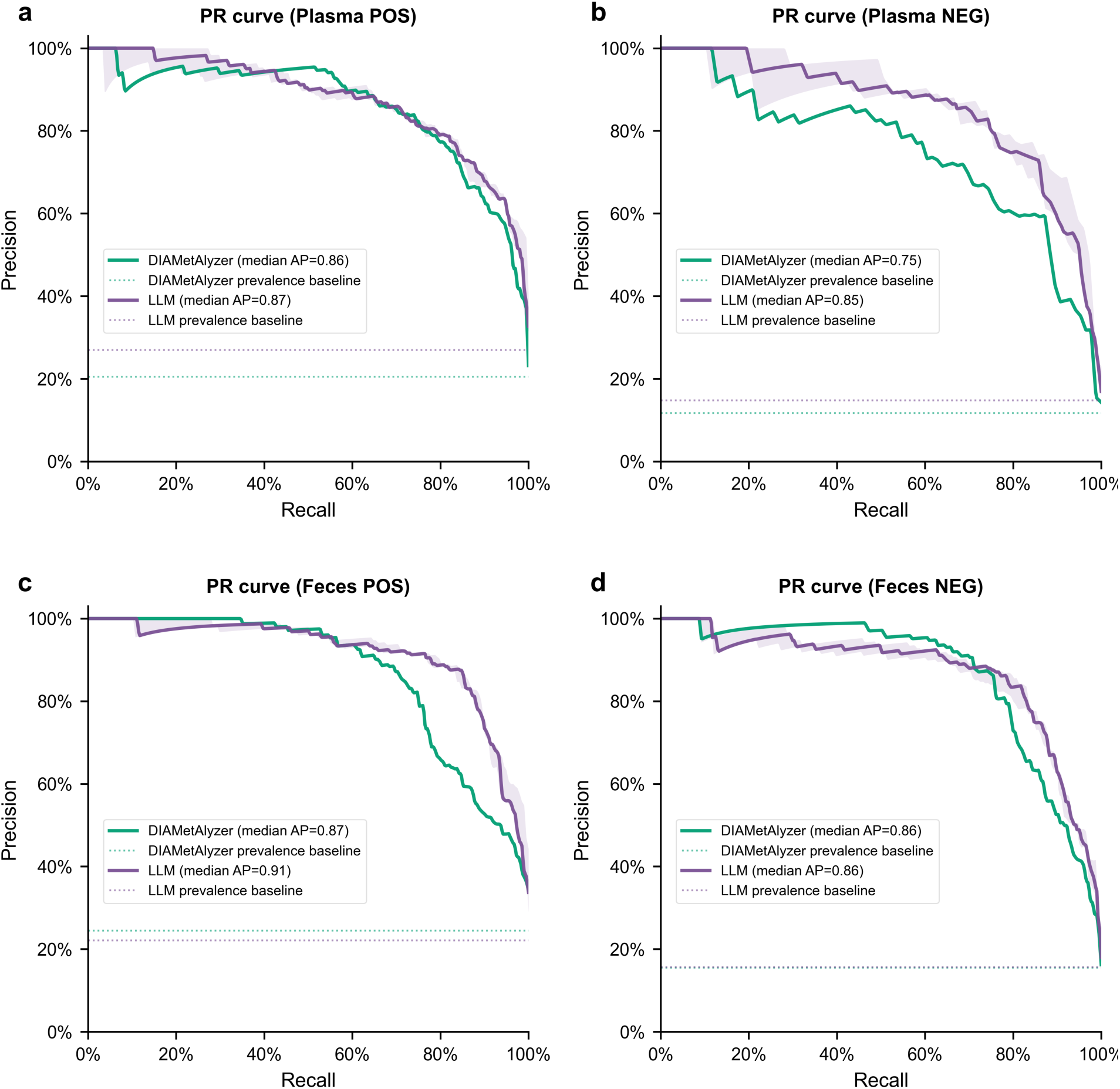
Within-dataset precision-recall performance of supervised candidate classifiers. Precision-recall curves are shown for Plasma POS (a), Plasma NEG (b), Feces POS (c), and Feces NEG (d). Out-of-fold predicted probabilities for each candidate were averaged across the 10 cross-validation repeats before calculating precision-recall curves and average precision (AP). LLM curves show the median and range across R0-R2, and LLM legend AP values represent the median across these three runs. DIAMetAlyzer curves and AP values were calculated from its averaged out-of-fold predictions. Dotted horizontal lines indicate the candidate-prevalence baseline for each candidate set.

**Supplementary Figure 6.**
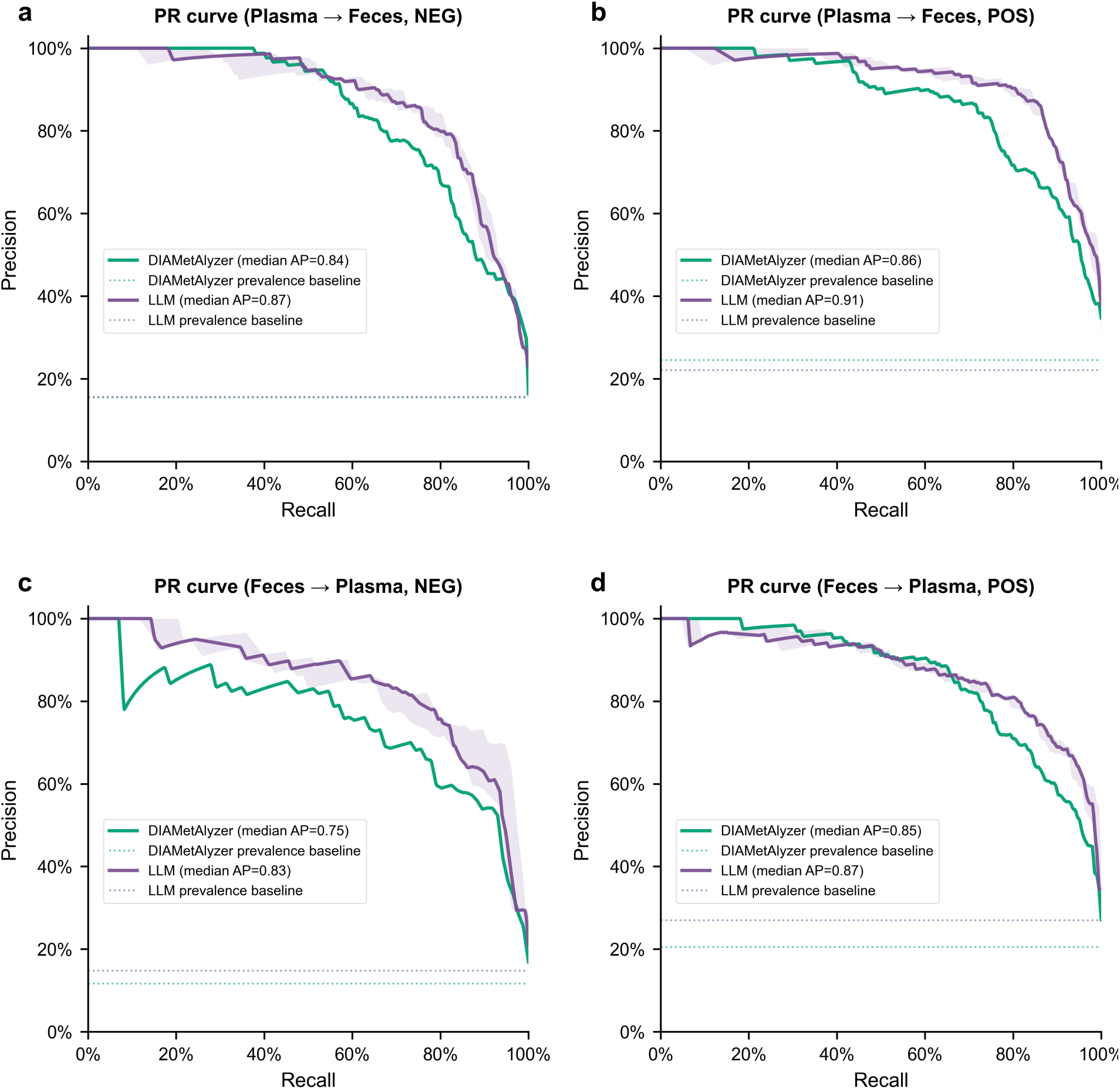
Precision-recall performance under bidirectional cross-dataset validation. Models trained on plasma were tested on feces in NEG (a) and POS (b) modes, whereas models trained on feces were tested on plasma in NEG (c) and POS (d) modes. Training and testing were restricted to matching ionization polarities. LLM curves show the median and range across matched R0-R2 models. Dotted lines indicate test-set candidate prevalence.

**Supplementary Table 1.** FDR-coverage of DIA lipidomics methods across plasma and fecal datasets. Coverage for OpenLipid LLM Majority, DIAMetAlyzer, and MS-DIAL DIA is expressed as the percentage of all manually annotated target lipids accepted at the indicated empirical FDR threshold.

| **Model** | **Plasma POS** | **Plasma NEG** | **Feces POS** | **Feces NEG** |
| --- | --- | --- | --- | --- |
| LLM Majority at 1% FDR | 35.7% | 12.2% | 39.2% | 22.7% |
| DIAMetAlyzer at 1% FDR | 33.7% | 4.8% | 70.7% | 15.5% |
| MS-DIAL DIA at 1% FDR | 6.0% | 0.0% | 16.6% | 0.0% |
| LLM Majority at 5% FDR | 55.3% | 19.0% | 71.8% | 31.8% |
| DIAMetAlyzer at 5% FDR | 57.8% | 19.7% | 89.0% | 17.8% |
| MS-DIAL DIA at 5% FDR | 12.6% | 0.0% | 47.5% | 0.0% |

**Supplementary Table 2.** Cross-dataset transfer of training-derived FDR thresholds. Probability cutoffs satisfying nominal FDR thresholds of 1% and 5% were determined exclusively from averaged out-of-fold predictions in the training dataset, using the highest-probability candidate for each target, and were applied unchanged to the external test dataset. Test FDR is the proportion of selected targets whose top-ranked candidate was incorrect. Test coverage is the number of selected targets divided by the total number of manually annotated targets in the test dataset, including both lipid-present and lipid-absent targets. LLM values are reported in R0/R1/R2 order.

| Direction | Polarity | Model | Training FDR | Fixed cutoff | Test FDR | Test coverage |
| --- | --- | --- | --- | --- | --- | --- |
| Plasma → Feces | NEG | DIAMetAlyzer | 1% | 0.980 | 0.00% | 10.6% |
| Plasma → Feces | NEG | DIAMetAlyzer | 5% | 0.980 | 0.00% | 10.6% |
| Plasma → Feces | NEG | LLM (R0/R1/R2) | 1% | 0.963 / 0.968 / 0.977 | 0.00% / 0.00% / 3.57% | 11.4% / 6.4% / 10.6% |
| Plasma → Feces | NEG | LLM (R0/R1/R2) | 5% | 0.955 / 0.968 / 0.954 | 2.86% / 0.00% / 3.64% | 13.3% / 6.4% / 20.8% |
| Plasma → Feces | POS | DIAMetAlyzer | 1% | 0.990 | 0.00% | 4.4% |
| Plasma → Feces | POS | DIAMetAlyzer | 5% | 0.920 | 7.89% | 63.0% |
| Plasma → Feces | POS | LLM (R0/R1/R2) | 1% | 0.974 / 0.984 / 0.964 | 0.00% / 0.00% / 1.67% | 27.6% / 7.2% / 33.1% |
| Plasma → Feces | POS | LLM (R0/R1/R2) | 5% | 0.962 / 0.978 / 0.952 | 4.44% / 2.38% / 1.16% | 49.7% / 23.2% / 47.5% |
| Feces → Plasma | NEG | DIAMetAlyzer | 1% | 0.995 | 14.29% | 4.8% |
| Feces → Plasma | NEG | DIAMetAlyzer | 5% | 0.897 | 27.59% | 59.2% |
| Feces → Plasma | NEG | LLM (R0/R1/R2) | 1% | 0.991 / 0.984 / 0.987 | 0.00% / 5.88% / 0.00% | 10.2% / 11.6% / 7.5% |
| Feces → Plasma | NEG | LLM (R0/R1/R2) | 5% | 0.985 / 0.959 / 0.977 | 6.67% / 14.29% / 10.53% | 20.4% / 38.1% / 25.9% |
| Feces → Plasma | POS | DIAMetAlyzer | 1% | 0.964 | 2.44% | 41.2% |
| Feces → Plasma | POS | DIAMetAlyzer | 5% | 0.899 | 7.91% | 69.8% |
| Feces → Plasma | POS | LLM (R0/R1/R2) | 1% | 0.986 / 0.987 / 0.988 | 2.00% / 9.09% / 3.23% | 25.1% / 5.5% / 15.6% |
| Feces → Plasma | POS | LLM (R0/R1/R2) | 5% | 0.969 / 0.959 / 0.970 | 7.89% / 7.21% / 8.26% | 57.3% / 55.8% / 54.8% |

